# Single-nucleus RNA-sequencing reveals novel potential mechanisms of ovarian insufficiency in 45,X Turner Syndrome

**DOI:** 10.1101/2023.12.22.572342

**Authors:** Sinéad M McGlacken-Byrne, Ignacio Del Valle, Theodoros Xenakis, Jenifer P Suntharalingham, Lydia Nel, Danielle Liptrot, Berta Crespo, Olumide K Ogunbiyi, Paola Niola, Tony Brooks, Nita Solanky, Gerard Conway, John C Achermann

**Author notes:** Corresponding author: Sinéad McGlacken-Byrne, Wellcome Trust Clinical Training Fellow Genetics and Genomic Medicine, UCL Great Ormond Street Institute of Child Health University College London, London, WC1N 1EH.

## Abstract

**Study question:** Can single-nuclei and bulk RNA sequencing technologies be used to elucidate novel mechanisms of ovarian insufficiency in Turner Syndrome (TS)?

**Summary answer:** Using single-nucleus and bulk RNA sequencing approaches, we identified novel potential pathogenic mechanisms underlying ovarian insufficiency in TS including and beyond X chromosome haploinsufficiency.

**What is known already:** Turner syndrome (TS) is the most common genetic cause of Primary Ovarian Insufficiency (POI) in humans. Morphological analyses of human fetal 45,X ovaries have demonstrated fewer germ cells and marked apoptosis established by 15-20 weeks post conception (wpc); however, we do not understand why POI develops mechanistically in the first instance.

**Study design, size, duration:** Single-nucleus RNA sequencing (snRNA-seq): two 46,XX and two 45,X (TS) human fetal ovaries at 12-13 wpc. Bulk RNA sequencing: 19 human fetal ovary, 20 fetal testis, and 8 fetal control tissue (n=47 total samples; Carnegie Stage 22-16wpc).

**Participants/materials, setting, methods:** To identify novel potential mechanisms of ovarian insufficiency in TS and to characterise X chromosome gene expression in the 45,X ovary, we performed snRNA-seq of peri-meiotic 46,XX (n=2) and 45,X (n=2) fetal ovaries at 12-13 weeks post conception (wpc); and 2) a bulk RNA sequencing time-series analysis of fetal ovary, testis, and control samples across four developmental timepoints.

**Main results and the role of chance:** Germ and somatic cell subpopulations were mostly shared across 46,XX and 45,X ovaries, aside from a 46XX-specific/45,X-depleted cluster of oogonia (“synaptic oogonia”) containing genes with functions relating to sex chromosome synapsis; histone modification; intracellular protein regulation and chaperone systems. snRNA-seq enabled accurate cell counting localised to individual cell clusters; the 45,X ovary has fewer germ cells than the 46,XX ovary in every germ cell subpopulation, confirmed by histopathological analysis. The normal sequence of X-chromosome inactivation and reactivation is disrupted in 45,X ovaries; *XIST* was not expressed in 45,X somatic cells but was present in germ cell clusters, albeit with lower expression than in corresponding 46,XX clusters. The 45,X ovary has a globally abnormal transcriptome, with low expression of genes with proteostasis functions (*RSP4X*); cell cycle progression (*BUB1B*); and OXPHOS mitochondrial energy production (*COX6C, ATP11C*). Genes with higher expression in 45,X cell populations were enriched for apoptotic functions (e.g., *NR4A1*).

**Limitations, reasons for caution:** Limitations include the relatively small sample size of the snRNA-seq analysis and the focus on a fixed meiotic timepoint which may overlook a dynamic process over time.

**Wider implications of the findings:** We characterise the human fetal peri-meiotic 45,X ovary at single-cell resolution and offer insights into novel pathogenic mechanisms underlying ovarian insufficiency in TS. Although asynapsis due to X chromosome haploinsufficiency likely plays a significant role, these data suggest meiotic failure and ovarian insufficiency may be a combinatorial process characterised by periods of vulnerability throughout early 45,X germ cell development

**Study funding/competing interest(s):** This research was funded in whole, or in part, by the Wellcome Trust Grants 216362/Z/19/Z to SMcG-B and 209328/Z/17/Z to JCA. Human fetal material was provided by the Joint MRC/Wellcome Trust (Grant MR/R006237/1) Human Developmental Biology Resource (http://www.hdbr.org). Research at UCL Great Ormond Street Institute of Child Health is supported by the National Institute for Health Research, Great Ormond Street Hospital Biomedical Research Centre (grant IS-BRC-1215-20012).

## Introduction

Turner syndrome (TS), the most common genetic cause of Primary Ovarian Insufficiency (POI) in humans, arises from a complete or partial loss of one X chromosome. Associated karyotypes include 45,X monosomy, 45,X mosaicism (45,X/46,XX or 45,X/46XY), isochromosome Xq (e.g., 46,X,i(Xq) or 45,X/46,X,i(Xq)), and ring X (e.g. 45,X/46,X,r(X)). Most women with TS have POI, although karyotypes with a greater X chromosome gene dosage (e.g,.45,X/46,XX) confer less of a POI risk (Cameron-Pimblett et al., 2017). However, even for women with 45,X monosomy, there is variability in the degree of ovarian function: while over 85% have disrupted pubertal progression and primary amenorrhoea, 15% present with secondary amenorrhoea or, occasionally, spontaneous pregnancy (Cameron-Pimblett et al., 2017).

Although TS is relatively common, occurring in 1/2000 female live births, and despite the high prevalence of POI in this condition, surprisingly few studies have investigated the pathogenesis and variability of ovarian dysfunction in TS. It is suggested that germ cell development follows a normal trajectory for the first trimester, followed by accelerated oocyte apoptosis and ovarian degeneration prior to birth (Singh & Carr, 1966; Hsueh et al., 1994; Sperling, 2008). Mouse studies from the 1980s demonstrated that 45,X ovaries are smaller and have fewer oocytes than 46,XX ovaries and proposed faulty sex chromosome pairing as the key driver of meiotic dysfunction (Burgoyne & Baker, 1985; Burgoyne & Baker, 1981). More recently, morphological analyses of human fetal 45,X ovaries demonstrated massive oocyte apoptosis by 15-20wpc; marked granulosa cell apoptosis; few or no viable follicles; and fewer oocytes compared to control samples (Lundgaard Riis et al., 2021; Modi et al., 2003; Reynaud et al., 2004). These data correlate with the clinical phenotype of POI in TS, but do not demonstrate mechanistically why POI develops in the first instance.

Considering the especial role sex chromosome genes play in mammalian gonadal development, and the importance of X-inactivation and reactivation to germ cell maturation, a reduction in X chromosome gene dosage due to X chromosome haploinsufficiency has been reasonably proposed to be central to the pathogenesis of ovarian insufficiency in TS (Afkhami et al., 2022; Heard & Turner, 2011; Severino et al., 2022). However, X chromosome gene dosage effects as the only contributing factor to ovarian dysfunction does not explain why some women with 45,X TS have relatively preserved reproductive function. Other possible explanations for the ovarian insufficiency phenotype include a reduction in dosage of genes escaping X chromosome inactivation (XCI escape genes); epigenetic abnormalities; downstream autosomal genetic disruption; and telomere length abnormalities (Trolle et al., 2016; Jackson-Cook, 2019; Rizzolio et al., 2007; Gravholt et al., 2022). Elucidating mechanisms of ovarian insufficiency in TS is important both for advancing our understanding of X chromosome biology as well as identifying novel therapeutic targets for this condition.

Here, we combine 1) a focused single-nucleus RNA-seq (snRNA-seq) analysis of peri-meiotic 46,XX (n=2) and 45,X (n=2) human fetal ovaries at 12/13 weeks post conception (wpc); 2) a bulk RNA sequencing (bulk RNA-seq) time-series analysis of human fetal ovaries extending from pre-meiosis (Carnegie Stage 22/23 (CS22/23)) until established meiosis (15/16wpc); and 3) histopathological analysis to investigate the role of X chromosome gene expression in human fetal ovary development and to reveal novel potential mechanisms of ovarian insufficiency in TS.

## Results

### Comparing the single cell landscapes of 46,XX and 45,X ovaries

snRNA-seq profiled 20,275 cells from four fetal ovaries (46,XX 12wpc (5802 cells), 45,X 12wpc (4570 cells), 46,XX 13wpc (4489 cells), and 45,X 13wpc (5414 cells)) to generate Uniform Manifold Approximation and Projection (UMAP) cell lineage projections (Supplementary Figure 1 and 2). Cell annotation was assigned based on the expression of established and recently identified markers and included germ cell clusters (in order of maturity: primordial germ cells (PGCs); fetal germ cells (FGCs); oogonia) and somatic cell clusters (gonadal (earlier) and ovarian (later) interstitial cells; ovarian surface epithelium cells (OSE); and early granulosa cells (pre-granulosa IIa (earlier) and IIb (later)) (Supplementary Table 1) (Garcia-Alonso et al., 2022; Gong et al., 2022). Data were then integrated and a UMAP representing this critical peri-meiotic developmental stage generated (Figure 1a).

**Figure 1:**
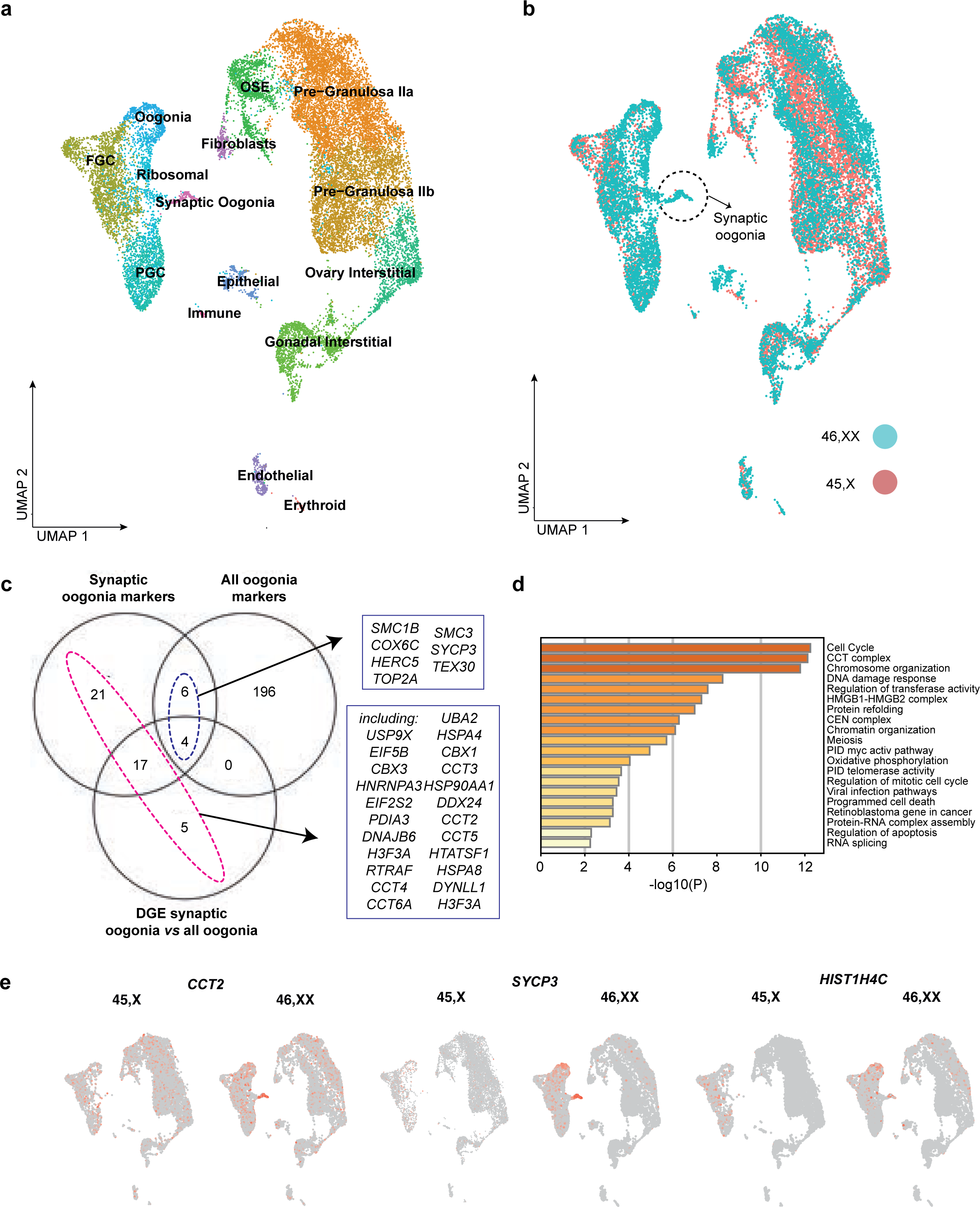
The peri-meiotic single-cell landscape of 45,X and 46,XX ovaries. **a)** UMAP (uniform manifold approximation and projection) of cell lineages across the integrated dataset, which represents 19,825 cells from four samples: 46,XX 12wpc (5765 cells), 45,X 12wpc (4302 cells), 46,XX 13wpc (4448 cells), and 45,X 13wpc (5321 cells). Individual cell populations are annotated. FGC, fetal germ cell; OSE, ovarian surface epithelium; PGC, primordial germ cell. Markers were based on prior knowledge and on previously published markers (Garcia-Alonso et al, *Nature,* 2022) **b)** The integrated UMAP split by karyotype. Germ and somatic cell subpopulations were mostly shared across 46,XX and 45,X ovaries. However, a 46,XX-specific population of “synaptic oogonia” were identified. **c)** Venn diagram demonstrating the top differentially expressed genes when comparing synaptic oogonia and all other clusters (n=48); the main oogonia cluster and all other clusters (n=206); and synaptic oogonia and the main oogonia cluster (n=28). Gene markers specific to the synaptic oogonia cluster (including genes differentially expressed in the synaptic oogonia cluster compared to the main oogonia cluster and/or genes differentially expressed in the synaptic oogonia cluster compared to all other clusters combined, and excluding genes also differentially expressed in the main oogonia cluster compared to other clusters) are highlighted in violet (n=43). Synaptic gene markers also differentially expressed in the main oogonia cluster are highlighted in pink (n=10). **d)** Gene enrichment analysis of the 53 genes positively differentially expressed in the 46,XX synaptic oogonia cluster compared to the main oogonia cluster and/or compared to all other clusters combined. **e)** UMAPs demonstrating differential expression and localisation of selected key synaptic oogonia markers.

Germ and somatic cell clusters were mostly shared across 46,XX and 45,X ovaries. Notably, there was one germ cell cluster with only three 45,X cells compared to 155 46,XX cells– essentially, a 46,XX-specific/45,X-depleted cluster (Figure 1b; Supplementary Table 2). To determine what set this population apart from other oogonia populations, differential gene expression analysis was performed between the 46,XX-specific oogonia population and the main oogonia population (log_2_FC>0.5, p.adj<0.05; 28 positively differentially expressed genes (DEGs) in the 46,XX-specific oogonia population) and between the 46,XX-specific oogonia population and all other clusters combined (pct.1 >0.6 (pct.1, percentage of cells expressing the gene in cluster 1), log_2_FC>0.5, p.adj<0.05; 48 positively DEGs in 46,XX-specific oogonia) (Figure 1c, Supplementary Tables 3 and 4). The combined list of 46,XX-specific oogonia markers (total of 53 genes) included genes relating to the meiotic cytoskeleton and synaptonemal complex, including *SYCP3, SMC1B,* and *SMC3* (Figure 1c; Supplementary Table 5). Gene enrichment analysis of genes expressed within this cluster confirmed an enrichment of terms related to these roles (Figure 1d). In addition to genes related to meiotic synapsis, genes relating to the broader peri-synapsis cytoskeleton were also highly expressed within the 46,XX-specific oogonia population (Figure 1e, Supplementary Figure 3). These included *DYNLL1,* which uses ATP hydrolysis to move nuclear-envelope tethered chromosomes along microtubules to facilitate homolog pairing in concert with the nucleoskeleton and cytoskeleton complex of SUN1 and KASH5 (Agrawal et al., 2022; Ding et al., 2007). Other highly expressed genes within the synaptic oogonia cluster included *RTRAF* (an RNA binding protein), *HNRNPA3* (a nuclear ribonucleoprotein), *HTATSF* (a transcription factor), and *HMGB2* (a chromatin protein), all with postulated roles in cytoskeleton assembly at meiosis (Ronfani et al., 2001). Highly expressed too within this cluster were genes playing central roles in molecular chaperone systems, including chaperonin containing TCP1 (CCT) complex genes (*CCT2, CCT3, CCT4, CCT5,* and *CCT6A*) and genes from heat shock protein chaperone systems (*HSP90AA1, HSPD1, DNAJA1, DNAJB6, PDIA3,* and *HSPA8)* (Figure 1e, Supplementary Figure 3). Taken together, the 46,XX-specific oogonia cluster was labelled ‘synaptic oogonia’.

### The 45,X ovary has fewer germ cells compared to 46,XX ovary

To examine cell populations *in vivo,* one 12wpc 45,X and one 12wpc 46,XX ovary were bisected and H&E stained (Figure 2a). The 45,X ovary was grossly smaller than the 46,XX ovary. Follicles and germ cells were visible in the 45,X ovary but were markedly fewer than in the 46,XX ovary, similar to what has been demonstrated using OCT4+ immunostaining previously (Lundgaard Riis et al., 2021).

**Figure 2:**
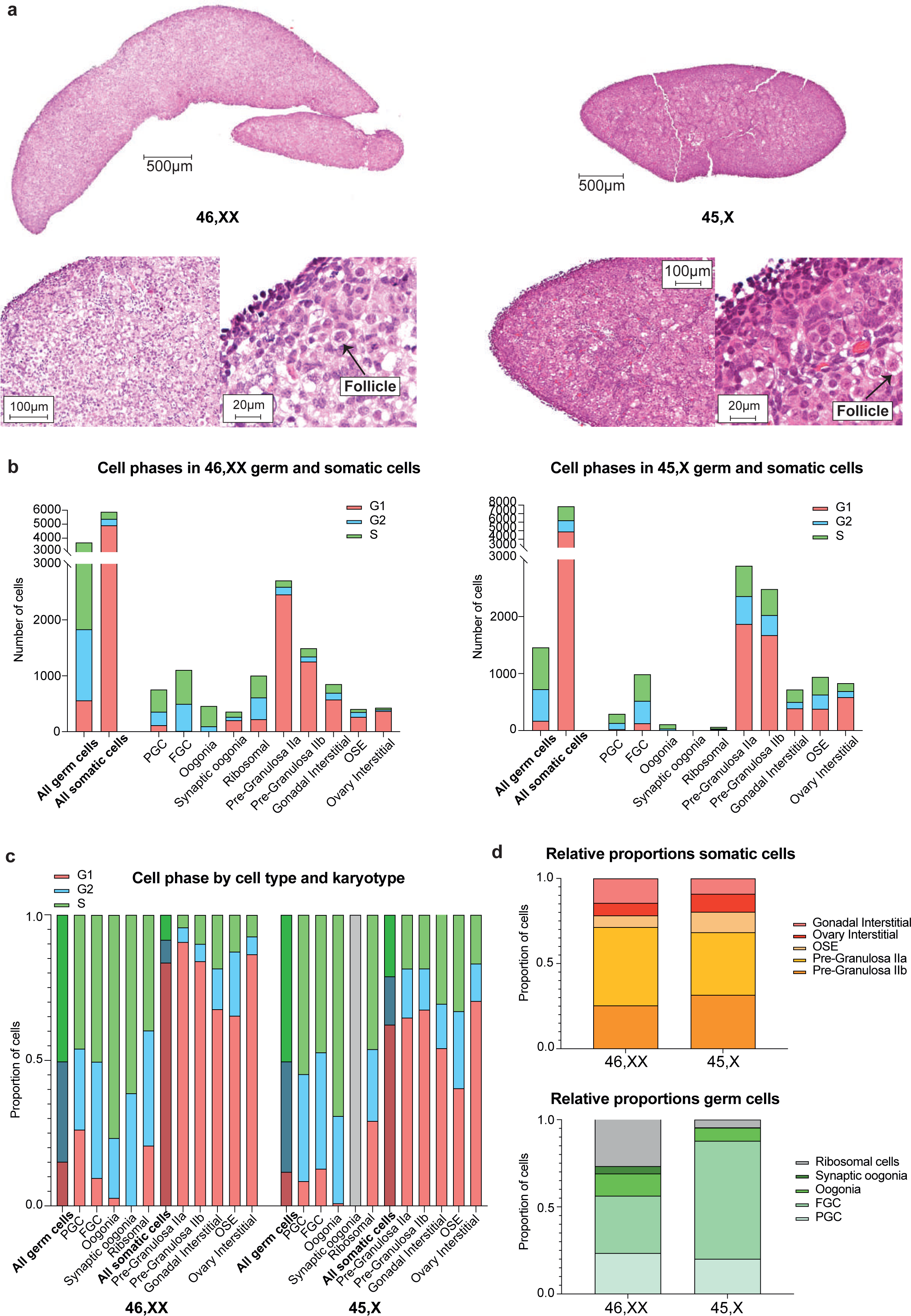
Comparing cell subpopulations between 46,XX and 45,X ovaries. **a)** Hematoxylin and eosin (H&E) staining of a 46,XX and 45,X 12wpc ovary. Arrow indicates an ovarian follicle**. b)** Total number of germ cells and somatic cells in the two 46,XX ovaries combined (left panel; total germ cells 3686, total non-germ cells 5882) and two 45,X ovaries combined (right panel; total germ cells 1489, total non-germ cells 7865), Selected subpopulations are shown based on annotation. The cell cycle status (G1, G2, or S) of all populations is indicated. **c)** For both 46,XX and 45,X ovaries, the proportion of cells in each subpopulation in G1, G2, or S cell cycle phase are shown**. d)** Relative proportions of somatic cell subpopulations (upper panel) and germ cell subpopulations (lower panel) for both 46,XX and 45,X ovaries.

Next, we analysed snRNA-seq data in detail to accurately compare and quantitate cell counts between 46,XX and 45,X ovaries within individual subpopulations and revealed that the 45,X ovary has fewer cells than the 46,XX ovary in every germ cell subpopulation (*p*<0.05; Figure 2b, Supplementary Table 2). Primordial germ cells (PGCs), fetal germ cells (FGCs), and oogonia within the 45,X ovary displayed similar proportions of cycling cells (i.e., in G2/S) compared to the 46,XX ovary (Figure 2c, Supplementary Table 2). There were proportionately more FGCs and fewer oogonia in 45,X ovaries compared to 46,XX, while relative proportions of somatic cells and PGCs were similar (p<0.05; Figure 2d; Supplementary Table 2).

### Distinct subsets of X chromosome genes are differentially expressed in the normal fetal 46,XX ovary

As ovarian insufficiency in TS is a potential X chromosome haploinsufficiency phenotype, we next characterised X chromosome gene expression in normal 46,XX human fetal ovary tissue using a bulk RNA-seq time-series analysis of 47 embryonic/fetal organs at four developmental stages (CS22-23 (7.5 to 8 wpc); 9-10wpc; 11-12wpc; 15-16wpc). This approach was used to identify X chromosome genes that are highly differentially expressed across this critical stage of development, which encompasses meiosis. This sample set included 19 ovaries (46,XX; n=5 biological replicates at each of CS22/23, 9/10wpc, and 11/12wpc; n=4 replicates at 15/16wpc); 20 testes (46,XY; n=5 replicates per developmental stage); and eight 46,XX control tissues (two different tissue samples per development stage, including spleen, skin, kidney, muscle, stomach, lung, and pancreas, balanced for a mix across endo-, meso-, and ectoderm cell lineages) (Supplementary Figure 4). Differential gene expression analyses of ovary compared to testes samples and to control samples were performed to identify groups of differentially expressed X chromosome genes (log_2_FC>2, p.adj<0.05; Supplementary Tables 6 and 7). Individual differential X chromosome gene expression analyses between ovary and testes samples were also conducted at four developmental timepoints (log_2_FC>2, p.adj<0.05; CS22/23, 9/10wpc, 11/12wpc, and 15/16wpc; Supplementary Tables 8-11).

Globally, there was an enrichment of positively differentially expressed X chromosome genes in the gonads: on ovary *v* testes and testes *v* ovary differential gene expression analyses, there were higher proportions of differentially expressed X chromosome genes compared to expected proportions (3.69%, or 894, of total coding genes are X chromosome genes; at log_2_FC>2, 6.01% of differentially expressed genes in the ovary across all stages were X chromosome genes (*p*<0.05)) (Supplementary Table 12). Individual sub-analyses at CS22/23, 9/10wpc, 11/12wpc, and 15/16wpc demonstrated that this X chromosome gene enrichment was maintained at earlier stages in the ovary (CS22-10wpc) but no longer present at 15/16wpc (Supplementary Table 12). Next, differentially expressed X chromosome genesets in the ovary compared to control samples and in the ovary compared to testis samples (log_2_FC>2, p.adj<0.05) were conflated to yield subsets of ovary-specific differentially expressed X chromosome genes (Figure 3a and 3b; Supplementary Figure 3), several of which have postulated roles in meiotic cytoskeleton organisation. *FAM9C* has an anti-apoptotic role in the testis and has homology to *SYCP3*, a known synaptonemal complex gene (Zhou et al., 2013). While *FAM9C* has not been associated with ovary development previously, the co-localisation of an X-linked gene within the same family, *FAM9B*, with *SYCP3* in human ovarian follicles at sex chromosome synapsis has been demonstrated (Yang et al., 2008). Here, expression of *FAM9C* localised to oogonia and increased at meiosis (Figure 3c; Supplementary Figure 3). *TEX11* is another gene of interest, where pathogenic variants are associated with sex chromosome asynapsis, meiotic arrest, and azoospermia in men; expression also localised to germ cell populations and also increased at meiosis in our data (Figure 3d) (Yatsenko et al., 2015). A similar expression pattern was seen for *BEND2*, a known X chromosome regulator of the peri-meiotic chromatic state in male mice (Figure 3e) (Ma et al., 2022). A previously published reproductive cell atlas (https://www.reproductivecellatlas.org) was used to validate the localised expression of these X chromosome genes (Supplementary Figure 3) (Vento-Tormo et al., 2018).

**Figure 3:**
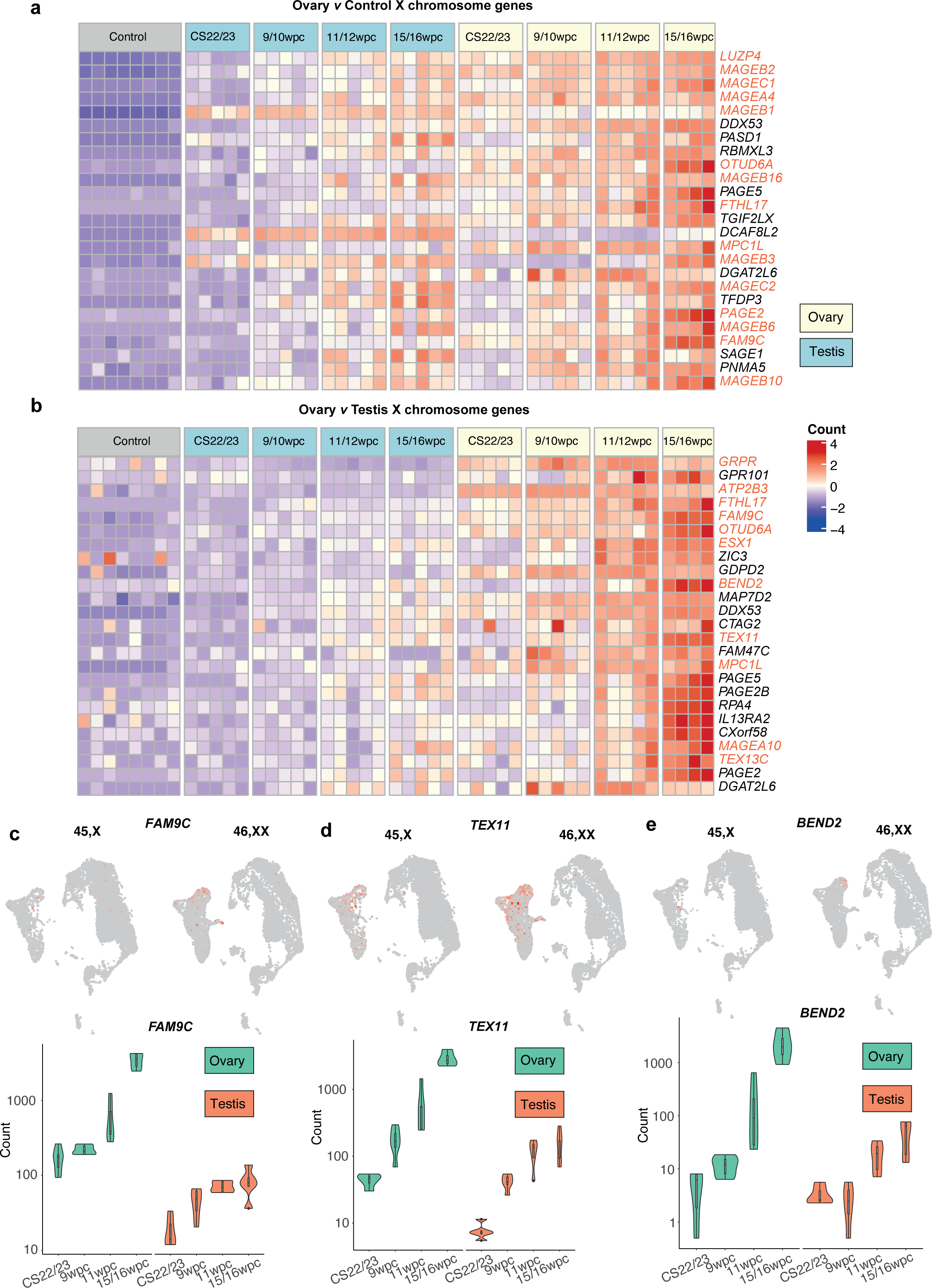
Differentially expressed X chromosome genes in the fetal ovary. **a)** Heatmap showing normalised gene expression values for the top 25 differentially expressed X chromosome genes in the ovary compared to control tissues displayed across the count matrix ordered according to descending log_2_FC values. A colour scale represents gene expression intensity (violet, lowest; red, highest). Genes discussed in the text are highlighted in orange. **b)** Heatmap showing normalised gene expression values for the top 25 differentially expressed X chromosome genes in the ovary compared to testis (p.adj<0.05). **c)-e)** Expression of *FAM9C, TEX11, BEND2*. *Upper panels:* UMAPs demonstrating increased expression of gene in oogonia populations in 46,XX compared to 45,X ovaries. *Lower panels:* Bulk RNAseq data demonstrating higher expression of these genes at 46,XX meiosis (15/16wpc) compared to at earlier stages.

As 46,XX meiosis involves a unique pattern of X-inactivation of one X chromosome within PGCs followed by X-reactivation in meiotic germ cells, we next performed a focused differential gene expression analysis of X chromosome genes known to escape X-chromosome inactivation (XCI escape genes). XCI escape genes were collated by merging lists from two recent studies identifying XCI escape genes (Supplementary Table 13) (Tukiainen et al., 2017; Wainer Katsir & Linial, 2019). There was an enrichment of XCI escape genes in the fetal ovary compared to testis (18.1% of differentially expressed genes in all ovary samples were XCI escape genes compared to an expected proportion of 13.6% (log_2_FC>0.5, p.adj<0.05) and there were more differentially expressed XCI genes in the ovary compared to testes at CS22/23, 9/10wpc, and 15/16wpc (p<0.05)) (Supplementary Table 14). Overlapping DEGs from the ovary *v* control and ovary *v* testis differential gene expression analyses demonstrated several highly ovary-specific XCI escape genes (log_2_FC>2, p.adj<0.05), including the above-discussed X chromosome genes *MPC1L*, *FAM9C*, *TEX11*, and *FAM9B,* as well as *KDM6A*, a histone demethylase (Figure 4a and 4b; Supplementary Figure 3, Supplementary Tables 6 and 7).

**Figure 4:**
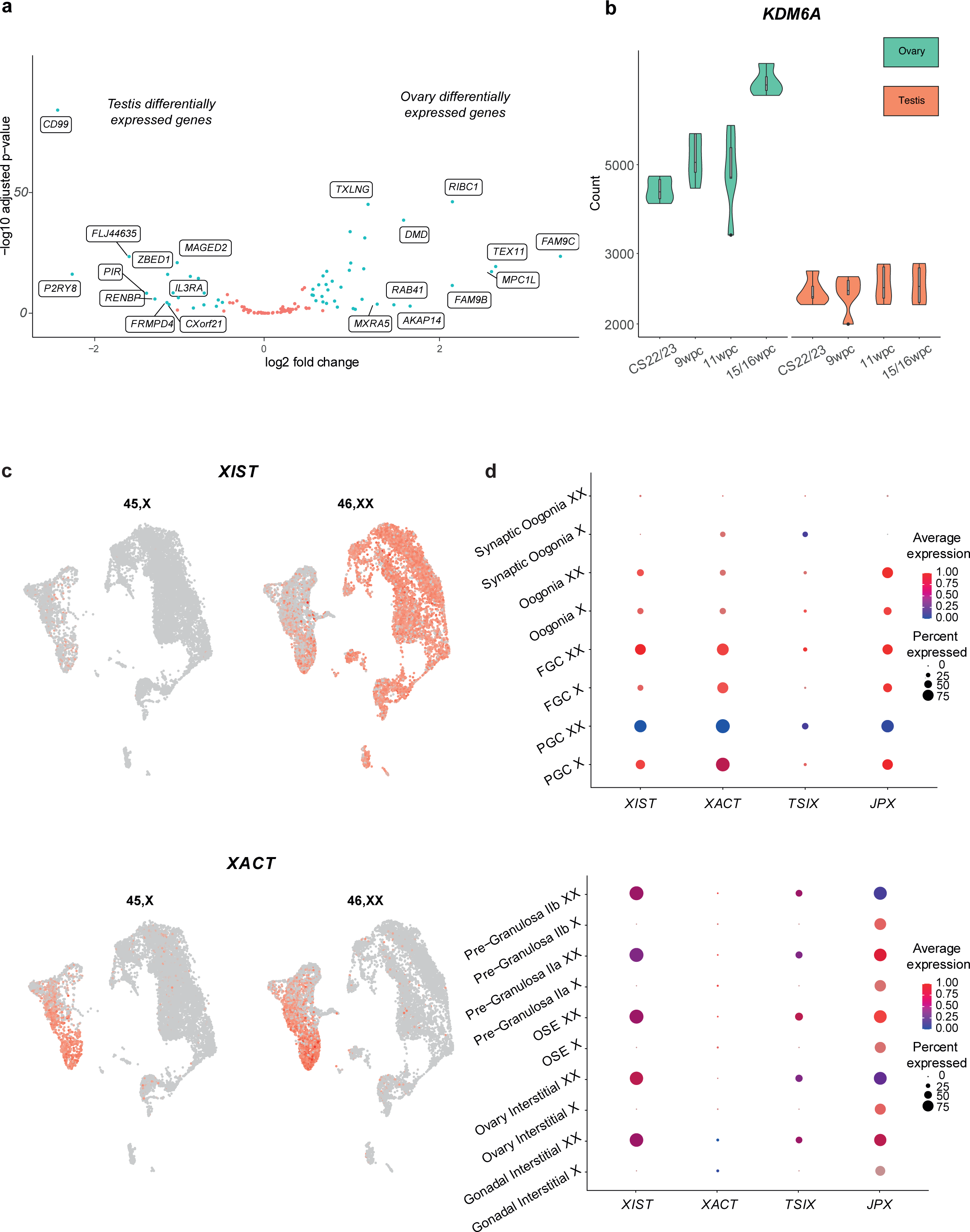
X-inactivation and reactivation patterns in 46,XX and 45,X ovaries. **a)** Volcano plot demonstrating differential gene expression analyses comparing all ovary *v* testis samples. Each dot represents a single gene. Genes that were differentially expressed with a log_2_FC>2 and p.adj<0.05 are coloured green; those coloured red did not meet these significance thresholds. Gene names of the top ten positively and negatively DEGs in this analysis are displayed. **b)** Bulk RNAseq data of *KDM6A* in the ovary and testis, demonstrating higher expression of *KDM6A* in the ovary compared to the testis and at 15/16wpc specifically. Note the non-linear scale. **c)** UMAPs demonstrating expression of *XIST* and *XACT* in individual subpopulations in the developing ovary. **d)** Dot plots showing the relative average expression and percentage of germ and somatic cell clusters expressing *XIST, XACT, TSIX,* and *JPX* in 46,XX and 45,X fetal ovaries. FGC, fetal germ cell; OSE, ovarian surface epithelium; PGC, primordial germ cell.

### X chromosome inactivation dynamics are disrupted in 45,X ovaries

To assess the contribution of XCI gene dosage to the transcriptomic landscape of 45,X meiotic ovaries, we compared expression of RNA genes involved in X-inactivation between 46,XX and 45,X ovaries. The 46,XX ovary expressed *XIST* in PGCs and FGCs, consistent with expected X-inactivation in early germ cells (Figure 4c and d). *XACT* expression, a key early step in X-reactivation required for germ cell development, was expressed in both PGCs and FGCs Peri-meiotic oogonia had low levels of *XACT* and *XIST,* suggesting X-inactivation was already erased in these cells. *JPX* was highly expressed in PGCs, FGCs, and oogonia. Synaptic oogonia had essentially no *XIST, JPX, TSIX,* or *XACT* expression as the X-inactivation process had by then been erased. *XIST* and *JPX* were expressed ubiquitously in somatic cells of the 46,XX ovary. Taken together, these data follow the predicted sequence of X-inactivation and reactivation in 46,XX oocytes. In contrast, however, substantial differences in X chromosome inactivation dynamics were seen in the 45,X ovary. *XIST* was not expressed in 45,X somatic cells but was present in PGCs, FGCs and oogonia, albeit at relatively lower percent expression levels than in the corresponding 46,XX populations. *JPX* was also expressed in these 45,X populations. Notably, *XACT* was expressed from the active X chromosome in 45,X PGCs, FGCs, and oogonia.

### The 45,X ovary has a globally abnormal transcriptome from the primordial germ cell stage

Next, differential gene expression analyses for each snRNA-seq subpopulation was performed to identify transcriptomic differences between 45,X and 46,XX ovaries (Figure 5a, Supplementary Tables 15-23).

**Figure 5:**
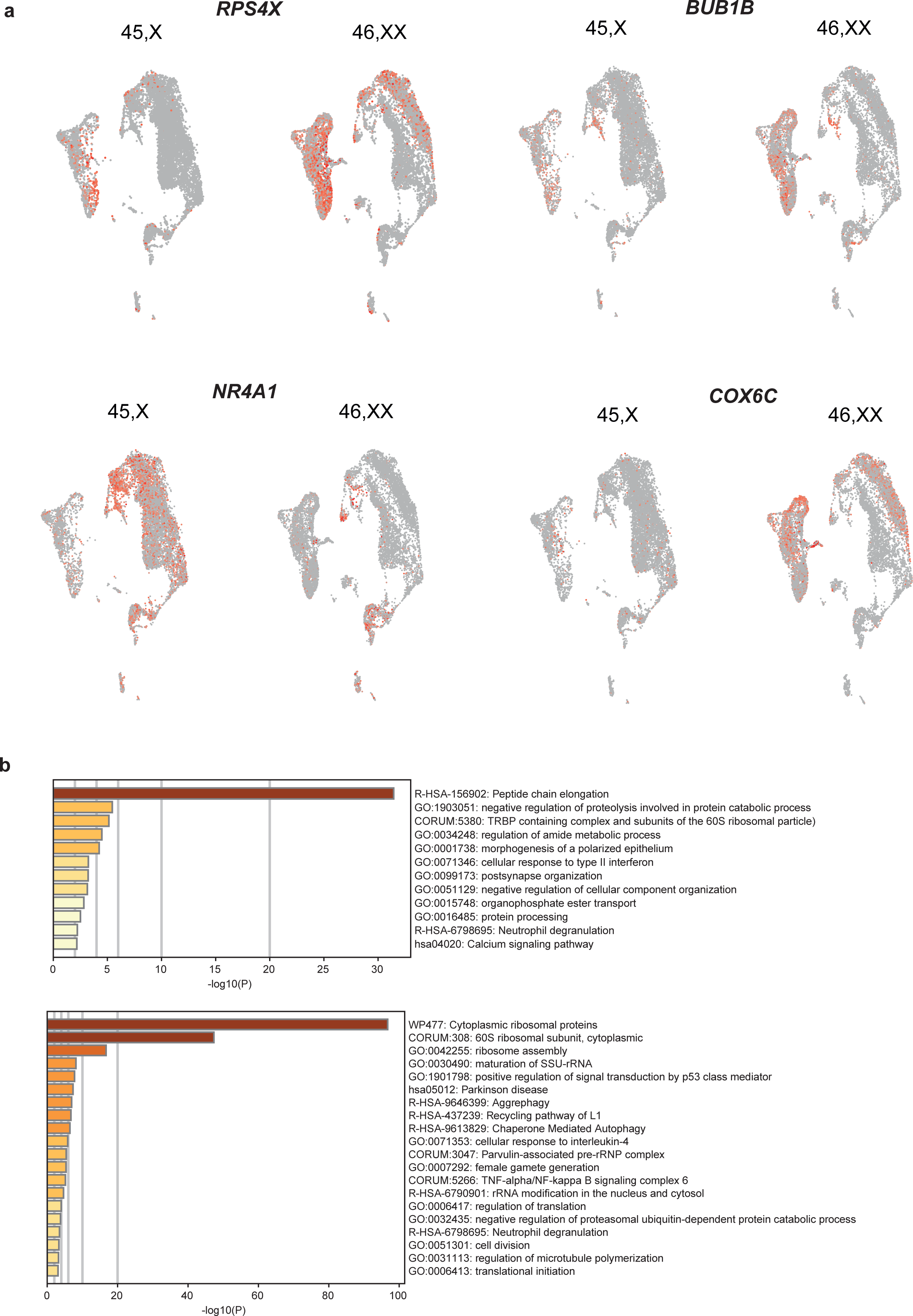
Differentially expressed genes between 46,XX and 45,X ovaries within individual cell clusters. **a)** UMAPs demonstrating differential expression and localisation of selected key differentially expressed genes. **b)** Gene enrichment analysis (Metascape) of genes positively differentially expressed in the 46,XX primordial germ cell (PGC) cluster compared to the 45,X PGC cluster (upper panel) and genes positively differentially expressed in the 46,XX fetal germ cell (FGC) cluster compared to the 45,X FGC cluster (lower panel).

On pathway enrichment analysis, at the PGC stage, genes involved in intracellular protein regulation (“proteostasis”) were expressed at a significantly lower level in the 45,X ovary. In the PGC and FGC clusters, DEGs (log_2_FC>0.5, p.adj<0.05) in 46,XX compared to 45,X PGCs were enriched for terms relating to proteostasis, including peptide chain elongation and proteolysis regulation (Supplementary Figure 5). These genes included *HUWE1,* an X-chromosome gene and E3 ubiquitin ligase (Moortgat et al., 2018); *EEF2,* required for GTP-dependent protein translation and cell proliferation (de Gassart et al., 2016); and *RPS4X,* an XCI escape gene required for ribosome biosynthesis (Figure 5a, Supplementary Table 15 and 24-27).

Additionally, several genes showed lower expression in the 45,X FGC populations compared to corresponding 46,XX FGC populations had functions relating to intracellular stress responses and epigenetic regulation. These genes included heat shock genes (*HSP90AB1, HSP90AA1, HSPA1A*), histone demethylases (*KDM1A*, *KDM3A*), and histone modifiers (*HIST1H2AA, HIST1H4C)*. This pattern extended to somatic pre-granulosa and interstitial cell populations (*HIST1H2AC* and *KDM3A*; Supplementary Tables 15 and 20-23).

Many genes with lower differential gene expression in 45,X oogonia compared to 46,XX oogonia had functions related to cell cycle progression (e.g., *BUB1, BUB1B*); meiotic chromosome organisation (*SYCP1, SYCP2, SYCP3*); and gametogenesis (Supplementary Figure 4 and 5; Supplementary Tables 4 and 15). Genes related to meiotic chromosomal organisation were also more lowly expressed in earlier PGC 45,X populations compared to 46,XX, including *DOCK11,* a Rho GTPase that facilitates actin cytoskeletal assembly required for cell migration (Qadir et al., 2015), and *RPS3,* a ribosomal gene which regulates microtubule polymerisation, spindle formation, and chromosome movement (Jang et al., 2012). There was also lower expression of ribosomal genes within the 45,X oogonia population compared to 46,XX oogonia which related both to proteostasis and to OXPHOS mitochondrial energy production, including *COX6C, ATP11C,* and *ATP5F1E*.

Not all differentially expressed genes were lower in 45,X clusters. A subset of genes had higher differential expression in 45,X clusters compared to 46,XX clusters, particularly within somatic cell populations. These genes were in pathways that were enriched for functions related to cell death and apoptosis, including *DUSP1*, a driver of apoptosis within granulosa cells (Yang et al., 2019); and *NR4A1* (also known as *NUR77/NGFIB*), a nuclear receptor expressed in the ovary and adrenal gland and involved in cell cycle regulation, steroidogenesis, inflammation, autophagy and apoptosis (Figure 5a-b; Supplementary Figure 5; Supplementary Tables 15 and 19-23) (Achermann et al., 2017). Interestingly, anti-apoptotic markers were also expressed in 45,X populations: *YBX3, MCL1, STRADB, XIAP,* and *BCL2* (Supplementary Figure 5; Supplementary Tables 15-23).

## Discussion

Ovarian insufficiency is a very common clinical manifestation of TS and understanding its natural history and the mechanisms driving it are important for reproductive counselling and decisions around fertility preservation in TS. For example, ovarian cryopreservation in young children with TS is being trialled in some countries as part of research studies (e.g., The Danish TURNER Cryopreservation Study, NCT05740579; NIH Gonadal Tissue Freezing for Fertility Preservation in Individuals at Risk for Ovarian Dysfunction, NCT04948658). Particularly, establishing the degree to which ovarian insufficiency in TS is established by the time of birth is a key consideration when planning therapeutic interventions.

Single-cell-omics technologies have been used to generate large tissue-specific “atlases”, such as a recent atlas of human fetal ovary development, which are very useful in understanding mechanisms of normal organogenesis (Vento-Tormo et al., 2018). However, by comparing unaffected control tissue with tissue affected by a condition, single-cell sequencing can also be used to discern the genetic drivers underlying pathogenic mechanisms. This work demonstrates the value of single-nuclei/single-cellRNA-sequencing technologies when applied to a developmental phenotype – in this case, TS. Specifically, we advance our understanding of X chromosome gene expression in human fetal ovary development, describe important transcriptomic differences between 46,XX ovaries and 45,X fetal ovaries at a critical peri-meiotic timepoint, and provide insight into novel potential mechanisms of ovarian insufficiency in TS.

Previous work has demonstrated that fetal ovaries in TS are smaller, have fewer follicles, and undergo marked cellular apoptosis during fetal life (Lundgaard Riis et al., 2021; Modi et al., 2003; Reynaud et al., 2004). These studies relied on 2D histological analysis of fetal ovary tissue in TS and used specific cell markers (e.g., OCT3/4) to estimate numbers of cells within the tissue overall. Here, we also demonstrate that 45,X fetal ovaries are macroscopically smaller with fewer follicles on histological analysis. However, for the first time to our knowledge, we use single-nucleus sequencing to accurately quantify cell counts within defined subpopulations of cells within TS ovaries compared to 46,XX ovaries. This unequivocally demonstrates that the 45,X ovary has fewer germ cells compared to 46,XX ovaries from the earliest point in primordial germ cell development.

We propose that the 46,XX-specific synaptic oogonia population identified on snRNA-seq analysis likely represents biological activity that does not occur in 45,X oogonia. An explanation is that asynapsis and broader peri-synapsis cytoskeleton dysfunction independently contribute to ovarian insufficiency in TS. This may be due to haploinsufficiency of key X-chromosome genes required for synapsis; for example, *HNRNPA3,* highly expressed in synaptic oogonia, interacts with XCI escape and synaptonemal complex gene *SMC1A;* and *HTATSF1,* another highly expressed gene within synaptic oogonia, itself escapes X-inactivation (Trolle et al., 2016; Severino et al., 2022). However, peri-meiotic X chromosome haploinsufficiency as the only mechanism underlying ovarian insufficiency in TS does not explain why there are fewer total germ cells in Turner syndrome *prior* to the point of meiotic synapsis; why there are markers of apoptosis from very early on in 45,X ovaries; why there is a dysregulated transcriptome throughout 45,X fetal ovary development rather than specifically at meiosis; and why some women with 45,X TS have a relatively preserved germ cell pool. Therefore, it is possible that some mechanisms underlying ovarian insufficiency in TS are established prior to sex chromosome synapsis, and the 46,XX-specific synaptic oogonia population arises because 45,X oogonia fail to progress to that developmental stage given preceding aberrations in germ cell maturation.

Combining bulk RNAseq and snRNA-seq demonstrates that both X chromosome gene expression and X chromosome gene dosage play important roles in the typical developing ovary and may contribute to ovarian insufficiency in TS due to roles X chromosome genes have both before and during sex chromosome synapsis. For example, several X chromosome and XCI escape genes (e.g., *BEND2, FAM9C, TEX11*) with functions related to the meiotic cytoskeleton are expressed in the mature meiotic oogonia stage in 46,XX ovaries but less expressed in the 45,X ovaries. However, 45,X ovaries have globally disrupted X-inactivation patterns that may contribute to ovarian insufficiency beyond meiotic synapsis. The scRNAseq data recapitulated X-inactivation patterns expected in normal 46,XX germ cells, but showed that these patterns are disrupted in the 45,X ovary (Severino et al., 2022; Heard & Turner, 2011). The lack of *XIST* expression in somatic cells of 45,X ovaries is unsurprising: *XIST* equalizes X chromosome gene expression with that of a 46,XY male, and, with only one X chromosome, this is not required in 45,X cells. The expression of *XIST* from 45,X germ cells, albeit in fewer than expected, may relate to the especial role that *XIST* plays in germ cell development: PGCs that have never been X-inactivated, or have been reactivated too quickly, do not progress through meiosis normally and display an abnormal mitotic profile (Severino et al., 2022). Possibly, some 45,X PGCs express *XIST* in an effort to inactivate their single X chromosome. *XACT* expression in 45,X FGCs at comparable expression to 46,XX FGCs further suggests that 45,X germ cells potentially follow the X-inactivation followed by X-reactivation pattern required for a normal meiotic programme. The X chromosome contains genes required for fundamental mitochondrial processes and complete X-inactivation in the setting of X chromosome haploinsufficiency likely would result in cell death. The fate of 45,X germ cells expressing *XIST* is therefore unclear – perhaps *XIST* expression in scattered 45,X cells is a mechanism of germ cell apoptosis in itself; or perhaps *XIST* expression is transient, localized to only a segment of the X chromosome, or downregulated before germ cell viability is threatened.

Beyond X-chromosome haploinsufficiency, and the impact of abnormal X-inactivation patterns on this, our data suggest additional possible drivers of ovarian dysfunction in TS. This includes an apparent dysregulated methylation programme in 45,X ovaries. Histone demethylases *KDM6A, KDM3A,* and *KDM1A* were differentially expressed in both 46,XX somatic and germ cell populations compared to 45,X (Cui et al., 2018). Lower expression and hypomethylation of *KDM6A,* an XCI escape gene proposed to regulate reproductive-related genes in mice, have been described in the peripheral leucocytes of women with TS (Berletch et al., 2013; Trolle et al., 2016). In mice, *Kdm3a* requires Hsp90 to demethylate H3K9me2 and to facilitate actin/tubulin folding within the cytoplasmic structures of maturing spermatids(Kasioulis et al., 2014). In the context of oestrogen receptor signalling, *Kdm1a* promotes gene expression by demethylating H3K9me2 and specifically is required to correctly establish DNA methylation of key imprinted genes (Wang et al., 2009; Katz et al., 2009). The enrichment of H2A histone modifiers *HIST1H2AC, HIST1H2AA,* and *HIST1H4C* across 46,XX cell populations is further evidence for a dysregulated methylome. These proteins interact with histone H3 by recruiting KDM1A to demethylate H3K9me2 and interact with ESR1 to regulate apoptosis and to enhance oestrogen responsive signalling and expression of cell cycle progression genes (Doisneau-Sixou et al., 2003; Perillo et al., 2000; Perillo et al., 2008; Su et al., 2014).

We also propose disrupted proteostasis as a further novel potential mechanism underlying ovarian insufficiency in TS, possibly due to an oocyte-specific energy deficiency. Genes within the synaptic oogonia cluster had roles in molecular chaperone systems required for proteostasis, including the CCT complex and heat shock protein family (Kim et al., 2013). Protein production results in the release of three-dimensional protein structures into the cellular environment which are at risk of aberrant folding and toxic aggregation (Hartl et al., 2011). The CCT is ATP-dependent and folds proteins important for microtubule integrity. The heat shock proteins also facilitate efficient protein folding and suppress apoptosis from toxic protein aggregates, again often requiring ATP. The synaptic oogonia cluster also contained several genes interacting as part of the ubiquitin-proteasome pathway, including *EIF5* and *EIF2S2*, two translation initiating factors specifically governing 40S ribosomal activity(Llácer et al., 2018) and which have established roles in translational regulation during oogenesis in mice and *Drosophila* (Villaescusa et al., 2006; Nakamura et al., 2004). Genes with antagonistic roles in neddylation, SUMOylation, ISGylation, and ubiquitination were also represented with the synaptic cluster. These post-translational protein modifications regulate the balance between protein production and degradation. Genes with roles in proteostasis were positively differentially expressed in 46,XX ovaries from the earliest germ cell stages, including *HUWE1* (ubiquitination), *EEF2* (translation), heat shock genes (*HSP90AB1, HSP90AA1, HSPA1A)*; and genes relating to ribosomal biogenesis (*RPS3, RPS4X).* Particularly, the role of *RPS4X* haploinsufficiency in the TS phenotype has been considered previously and we suggest that *RPS4X* haploinsufficiency in the 45,X fetal ovary results in impaired ribosomal assembly and reduced protein synthesis (Peeters et al., 2019; Fisher et al., 1990). Protein synthesis demands high intracellular energy needs and several genes with proteostasis roles specifically require ATP to function. Indeed, there was an apparent deficiency in the expression of energy mitochondrial genes, such as *COX6C, ATP11C, ATP5F1*, in 45,X oogonia. Possibly, the apparent predominance of genes related to proteostasis within 46,XX oogonia reflects the inherent role of carefully controlled protein production during the energy-demanding processes of meiosis and sex chromosome synapsis which are impaired in 45,X ovaries due to issues arising earlier in germ cell development. However, an alternative explanation is that oogenesis, and particularly synapsis, involves high intracellular protein and energy production demands which 45,X oocytes cannot efficiently meet, possibly due to haploinsufficiency of genes required for proteostasis or even a primary energy deficiency in 45,X oogonia.

Limitations of this study include the relatively small sample size included in the snRNA-seq analysis (four in total) and the focus on a fixed timepoint in meiosis which may overlook a dynamic process over time and therefore miss important events occurring either pre-meiosis or once meiosis has been fully established. However, for the first time to our knowledge, we examine 45,X fetal tissue at single-cell resolution and offer insights into novel pathogenic mechanisms underlying ovarian insufficiency in TS. Taken together, our data suggest that, while X chromosome haploinsufficiency likely plays a significant role, a single, “all-or-nothing” event in the developing 45,X germ cell is unlikely to be solely responsible for meiotic failure and ovarian insufficiency in TS. Rather, we propose it may be a combinatorial process starting early in 45,X germ cell development characterised by periods of vulnerability throughout germ cell development.

## Methods

### Tissue samples

Human embryonic and fetal samples used for these studies were obtained from the Human Developmental Biology Resource (HDBR, a Medical Research Council (MRC) and Wellcome Trust-funded tissue bank regulated by the Human Tissue Authority (www.hdbr.org)). Appropriate maternal consent was obtained prior to sample collection. Full ethical approval was obtained from the NRES London-Fulham Ethics Committee (08/H0712/34+5, 18/LO/0822) and the Newcastle Ethics Committee (08/H0906/21+5, 18/NE/0290) with specific study approval from the HDBR (project numbers 200408; 200481; 200581). Ovaries, testes and 46,XX control tissues were visualised and isolated by blunt dissection. Embryonic and fetal tissue age were calculated by HDBR researchers in London and Newcastle using accepted staging guidelines, such as Carnegie staging for embryos up to 8wpc and foot length and knee-heel length in relation to standard growth data for older fetuses. Karyotyping by G-banding or quantitative PCR (chromosomes 13, 15, 16, 18, 21, 22, X, Y) were used to ascertain the sex of the embryo or fetus and to verify if the karyotype was 46,XX or 45,X. Samples were frozen at −70°C or stored in fixative (10% formalin or 4% paraformaldehyde) prior to use.

### Bulk RNA-sequencing

#### RNA extraction

RNA extraction was carried out utilising the AllPrep DNA/RNA Mini Kit from QIAGEN N.V. following the manufacturer’s protocol. Tissues preserved at −70 degrees were homogenized with an electronic pestle (Kimble, New Jersey). The minimum RNA quantity required for sequencing was 50ng with a 260/280 ratio >2.0. Additionally, RNA integrity was assessed by measuring the RNA Integrity Number (RIN) using an Agilent Bioanalyzer (Agilent, Santa Clara, CA). All samples exhibited a RIN value higher than 7.

#### Library preparation and RNA sequencing

The Hamilton StarLet (Hamilton, Reno, NV) robotic platform was used for library preparation and the Tapestation 4200 platform (Agilent, California, USA) was used for qualitative checks. Libraries were prepared using the KAPA RNA HyperPrep Kit followed by sequencing on the Illumina NovaSeq® (Illumina, San Diego, CA) at a minimum of 25 million paired end reads (75bp) per tissue sample.

#### Bioinformatic analysis

Quality control (QC) analysis of Fastq reads used the FastQC Subsequently, the reads were aligned to the GRCh38 genome (hg38) to generate BAM files using STAR 2.7 (Dobin et al., 2013). For gene expression quantification, differential gene expression analysis, and identification of expression patterns, featureCounts from the Subread package (v2.0.2) and DESeq2 (v1.28.1) were employed, respectively (Love et al., 2014; Liao et al., 2014).

During differential gene expression analysis, cut-off values of 0.05 for adjusted p-values and 1, 1.5, or 2 for log_2_fold changes were utilised. Gene annotation and pathway enrichment analysis were performed using Metascape (Zhou et al., 2019). Visualisation of differentially expressed genes was accomplished through heatmaps generated using the ComplexHeatmap package in R (Gu et al., 2016). Detailed information about the R packages and their versions utilised in the bioinformatic analysis pipeline can be found in Supplementary Methods 1.

### Single-nucleus RNA-sequencing (snRNA-seq)

#### Samples

Two 46,XX fetal ovaries (12wpc and 13wpc) and two 45,X ovaries (12wpc and 13wpc) were obtained from HDBR following ethical approval and consent as above. Tissue karyotype was confirmed on extracted DNA from skin biopsy and array. Samples were frozen at −70°C prior to use.

#### Single-nuclei dissociation

To prepare single-nuclei suspensions, a published protocol (Martelotto et al, Broad Institute, dx.doi.org/10.17504/protocols.io.bw6qphdw) was followed. Throughout the procedure, all samples and reagents were maintained on either wet ice or kept at 4°C. Tissue samples were diced into approximately 1mm³ pieces using a scalpel before dissociation in 300μL of Salty Ez10 Lysis Buffer supplemented with 0.2-0.5U/μL of RNase inhibitor (Sigma-Aldrich) using a 2ml Kimble douncer (Sigma-Aldrich; 10 strokes with a loose pestle and 10 strokes with a tight pestle).

An additional 700μL of chilled Salty Ez10 Lysis Buffer supplemented with 0.2-0.5U/μL of RNase inhibitor was added followed by gentle pipette mixing. The samples were then incubated on ice for five minutes, during which time they were gently pipette mixed 2-3 times. A MACS 70μm filter (Miltenyi Biotec) was placed in a 50ml Falcon tube kept on ice. The single-nuclei suspension was transferred to the filter, and the filtrate was collected in a 1.5ml LoBind tube (Eppendorf). The nuclei were centrifuged at four degrees at 500g for five minutes, and the supernatant was removed, leaving a pellet.

Salty Ez10 Lysis Buffer (1ml) was added to the pellet which was gently resuspended. The suspension was incubated on ice for an additional five minutes and centrifuged for five minutes as before. After removing the supernatant, 500μL of Wash and Resuspension Buffer 2 (WRB2) was added without disturbing the pellet. The sample was left on ice for five minutes before being gently resuspended. Cell counting was performed using the Luna-FL™ Dual Fluorescence Cell Counter (Logos Biosystems) and Acridine Orange/Propidium Iodide (AO/PI) Cell Viability Kit Counter (Logos Biosystems). Sample concentrations of cells per μl were: 46,XX: 12wpc 1720cells/μl; 45,X: 12wpc 1430cells/μl; 46,XX 13wpc: 1170cells/μl; 45,X 13wpc: 906cells/μl). Components for the two buffers, made up to a 100ml stock, are shown in Supplementary Methods 2.

#### Gel bead in EMulsion (GEM) Generation and Barcoding

Libraries from single nuclei suspensions were processed using the 10X Chromium Single Cell 3’ kit as per protocol (10X Genomics, Pleasonton, CA).

#### Single-nucleus sequencing

Single-nuclei libraries were pooled and sequenced on the Illumina NovaSeq S2 v1.5 platform. Sequencing was paired-end using single indexing and a minimum of 20,000 read pairs per cell.

#### Bioinformatic analysis

The R package Seurat (v4.0.2) was used to generate a single-cell matrix as described previously(Hao et al., 2021). Cycling cells were included. Quality control (QC) filtering retained cells with feature counts >400, percentage of mitochondrial genes <1%, and percentage of ribosomal genes <5% (Supplementary Figure 1).

SoupX (v1.6.2) was used to remove cell-free mRNA contamination, ParamSweep (v3.0) for parameter optimization, and doubletFinder (v2.0.3) to remove doublets. The count matrix was normalized and 3000 variable genes selected. After scaling, dimensionality reduction was performed using the first 30 principal components.

Seurat packages FindClusters and RunUMAP were used to identify cell clusters and for UMAP visualization. The clustree R package (v0.5) was used to select a clustering resolution of 0.3 throughout. SCTransform (v0.3.5) and FindIntegrationAnchors were used to integrate the 12wpc and 13wpc 46,XX ovaries. A mitochondrial cluster with high ribosomal content remained despite upstream QC; this cluster was removed from the initial analyses and the samples re-integrated to yield a final UMAP object and final cell counts. Differential gene expression was performed using the FindAllMarkers function (‘min.pct=0.25, logfc.threshold=0.25’). Functions within Seurat (FeaturePlot, VlnPlot, and DotPlot) were used to visualise gene marker expression. All R packages and versions used are listed in Supplementary Methods 1.

### Statistical Analysis

Statistical analyses were performed in R (v4.2.0). A *p* value of less than 0.05 was considered significant. The Benjamini-Hochberg approach was used to adjust for multiple testing with cut-off adjusted *p* values of 0.05 (Benjamini & Hochberg, 1995). Data are shown as individual data points or as violin plots as appropriate.

## Data availability and Supplementary data

### Data repository links

Bulk RNA sequencing data are deposited in ArrayExpress/Biostudies (accession number S-BSST693). Single-cell RNA sequencing data are deposited in ArrayExpress/Biostudies (accession number S-BSST1194)

## Supplementary Files

**Supplementary Materials:** Supplementary methods 1-2; Supplementary Figures 1-5

**Supplementary Table 1:** Gene markers used for annotation of cell clusters.

**Supplementary Table 2:** Comparing cell count per cluster between 45,X and 46,XX ovaries

**Supplementary Table 3:** Differential gene expression analysis of genes within the synaptic oogonia population compared to the main oogonia population.

**Supplementary Table 4:** Differential gene expression analysis of genes within the synaptic oogonia population compared to the main oogonia population.

**Supplementary Table 5:** Synaptic oogonia markers

**Supplementary Table 6:** Top 250 differentially expressed X chromosome genes, ovary v testis.

**Supplementary Table 7:** Top 250 differentially expressed X chromosome genes, ovary v control.

**Supplementary Table 8:** Differentially expressed X genes, ovary v testis CS22/23.

**Supplementary Table 9:** Top 250 differentially expressed X genes, ovary v testis 9/10wpc.

**Supplementary Table 10:** Differentially expressed X genes, ovary v testis 11/12wpc.

**Supplementary Table 11:** Differentially expressed X genes, ovary v testis 15/16wpc.

**Supplementary Table 12:** X chromosome gene enrichment in the developing gonad.

**Supplementary Table 13:** Genes escaping X chromosome inactivation.

**Supplementary Table 14:** Enrichment of XCI escape genes in the fetal ovary.

**Supplementary Table 15:** Key differentially expressed genes in 46,XX and 45,X ovaries across individual cell clusters.

**Supplementary Table 16:** 46,XX v 45,X differential gene expression analysis within the primordial germ cell cluster (PGC).

**Supplementary Table 17:** 46,XX v 45,X differential gene expression analysis within the fetal germ cell cluster (FGC).

**Supplementary Table 18:** 46,XX v 45,X differential gene expression analysis within oogonia.

**Supplementary Table 19:** 46,XX v 45,X differential gene expression analysis within ovarian surface epithelium (OSE).

**Supplementary Table 20:** 46,XX v 45,X differential gene expression analysis within gonadal interstitial cells.

**Supplementary Table 21:** 46,XX v 45,X differential gene expression analysis within ovary interstitial cells.

**Supplementary Table 22:** 46,XX v 45,X differential gene expression analysis within pre-granulosa IIA cells.

**Supplementary Table 23:** 46,XX v 45,X differential gene expression analysis within pre-granulosa IIB cells

## Author contributions

Author contributions were as follows: Study conceptualization: SMcG-B, IdV, JCA; Methodology: SMcG-B, IdV, TX, TB; Investigation: SMcG-B, IdV, TX, JPS, BC, OO, LN, DL, PN, TB, JCA; Formal analysis: SMcG-B, IdV; Data curation: SMcG-B, IdV; Resources: NS; Project administration: JCA; Supervision: GSC, NS, JCA; Validation: SMcG-B, IdV, TX, TB; Visualization: SMcG-B, IdV, JCA; Writing – original draft: SMcG-B, JCA; Writing – review & editing: All authors; Funding acquisition: SMcG-B, JCA.

## Acknowledgments

We thank other members of UCL Genomics and the Human Developmental Biology Resource for their additional contributions to this work. This work forms part of a PhD Thesis submitted to University College London (SMcG-B).

