## Supplemental Materials for "Single-nucleus RNA-sequencing reveals novel potential mechanisms of ovarian insufficiency in 45,X Turner Syndrome"

Running title: Novel ovarian insufficiency mechanisms in Turner Syndrome

Key words: Turner syndrome, single-nucleus sequencing, single-cell sequencing, ovarian insufficiency, X chromosome genetics

*Corresponding author:

Sinéad McGlacken-Byrne

Wellcome Trust Clinical Training Fellow

Genetics and Genomic Medicine

UCL Great Ormond Street Institute of Child Health

University College London

London, WC1N 1EH

#### Supplementary Methods 1:

R Packages used in scRNAseq analysis

| | | | |
|---|---|---|---|
| tidyr_1.2.1 | fansi_1.0.3 | listenv_0.9.0 | renv_0.16.0 |
| abind_1.4-5 | farver_2.1.1 | lmtest_0.9-40 | reshape2_1.4.4 |
| AnnotationDbi_1.56.2 | fastmap_1.1.0 | locfit_1.5-9.7 | reticulate_1.26 |
| AnnotationHub_3.2.2 | filelock_1.0.2 | magrittr_2.0.3 | rhdf5_2.38.1 |
| assertthat_0.2.1 | fitdistrplus_1.1-8 | MASS_7.3-58.1 | rhdf5filters_1.6.0 |
| beachmat_2.10.0 | future_1.30.0 | Matrix_1.5-3 | Rhdf5lib_1.16.0 |
| Biobase_2.54.0 | future.apply_1.10.0 | MatrixGenerics_1.6.0 | rlang_1.0.6 |
| BiocFileCache_2.2.1 | generics_0.1.3 | matrixStats_0.63.0 | ROCR_1.0-11 |
| BiocGenerics_0.40.0 | GenomeInfoDb_1.30.1 | memoise_2.0.1 | RSQLite_2.2.20 |
| BiocManager_1.30.19 | GenomeInfoDbData_1.2.7 | mime_0.12 | rstudioapi_0.14 |
| BiocParallel_1.28.3 | GenomicRanges_1.46.1 | miniUI_0.1.1.1 | Rtsne_0.16 |
| BiocVersion_3.14.0 | ggforce_0.4.1 | munsell_0.5.0 | S4Vectors_0.32.4 |
| Biostrings_2.62.0 | ggplot2_3.4.0 | nlme_3.1-161 | scales_1.2.1 |
| bit_4.0.5 | ggraph_2.1.0 | parallel_4.1.0 | scattermore_0.8 |
| bit64_4.0.5 | ggrepel_0.9.2 | parallelly_1.33.0 | sctransform_0.3.5 |
| bitops_1.0-7 | ggridges_0.5.4 | patchwork_1.1.2 | scuttle_1.4.0 |
| blob_1.2.3 | globals_0.16.2 | pbapply_1.6-0 | Seurat_4.3.0 |
| cachem_1.0.6 | glue_1.6.2 | pillar_1.8.1 | shiny_1.7.4 |
| celldex_1.4.0 | goftest_1.2-3 | pkgconfig_2.0.3 | SingleCellExperiment_1.16.0 |
| cli_3.5.0 | graphlayouts_0.8.4 | plotly_4.10.1 | sparseMatrixStats_1.6.0 |
| cluster_2.1.4 | grid_4.1.0 | plyr_1.8.8 | spatstat.data_3.0-0 |
| clustree_0.5.0 | gridExtra_2.3 | png_0.1-8 | spatstat.explore_3.0-5 |
| codetools_0.2-18 | gtable_0.3.1 | polyclip_1.10-4 | spatstat.geom_3.0-3 |
| colorspace_2.0-3 | HDF5Array_1.22.1 | progressr_0.12.0 | spatstat.random_3.0-1 |
| compiler_4.1.0 | htmltools_0.5.4 | promises_1.2.0.1 | spatstat.sparse_3.0-0 |
| cowplot_1.1.1 | htmlwidgets_1.6.0 | purrr_1.0.0 | spatstat.utils_3.0-1 |
| crayon_1.5.2 | httpuv_1.6.7 | R.methodsS3_1.8.2 | splines_4.1.0 |
| curl_4.3.3 | httr_1.4.4 | R.oo_1.25.0 | stringr_1.5.0 |
| data.table_1.14.6 | ica_1.0-3 | R.utils_2.12.2 | SummarizedExperiment_1.24.0 |
| DBI_1.1.3 | igraph_1.3.5 | R6_2.5.1 | survival_3.4-0 |
| dbplyr_2.2.1 | interactiveDisplayBase_1.32.0 | RANN_2.6.1 | tensor_1.5 |
| DelayedArray_0.20.0 | IRanges_2.28.0 | rappdirs_0.3.3 | tibble_3.1.8 |
| DelayedMatrixStats_1.16.0 | irlba_2.3.5.1 | RColorBrewer_1.1-3 | tidygraph_1.2.2 |
| deldir_1.0-6 | jsonlite_1.8.4 | R.oo_1.25.0 | tidyselect_1.2.0 |
| digest_0.6.31 | KEGGREST_1.34.0 | R.utils_2.12.2 | tools_4.1.0 |
| DoubletFinder_2.0.3 | KernSmooth_2.23-20 | R6_2.5.1 | tweenr_2.0.2 |
| dplyr_1.0.10 | labeling_0.4.2 | RANN_2.6.1 | utf8_1.2.2 |
| dqrng_0.3.0 | later_1.3.0 | rappdirs_0.3.3 | uwot_0.1.14 |
| DropletUtils_1.14.2 | lattice_0.20-45 | RColorBrewer_1.1-3 | vctrs_0.5.1 |
| edgeR_3.36.0 | lazyeval_0.2.2 | Rcpp_1.0.9 | viridis_0.6.2 |
| ellipsis_0.3.2 | leiden_0.4.3 | RcppAnnoy_0.0.20 | SoupX_1.6.2 |
| euratObject_4.1.3 | lifecycle_1.0.3 | RCurl_1.98-1.9 | ParamSweep_3.0 |
| ExperimentHub_2.2.1 | limma_3.50.3 | remotes_2.4.2 | SCTransform_0.3.5 |

**Supplementary Methods 2:**

Components for the buffers used for single-nuclei suspension:

*Salty Ez10 Lysis Buffer*

10 mM Tris-HCl pH 7.5 1M

146 mM NaCl 5M

1 mM $CaCl_2$ 1M

21 mM $MgCl_2$ 1M

0.03% Tween-20 (Sigma-Aldrich)

0.01% BSA (Miltenyi Biotec)

10% Ez Lysis Buffer (Sigma-Aldrich)

0.2-1 U/uL Protector RNAse Inhibitor (Roche)

1 mM DTT (ThermoFisher Scientific)

*Wash and Resuspension Buffer 2 (WRB2)*

10 mM Tris-HCl pH 7.5

10 mM NaCl

3 mM $MgCl_2$

1mM DTT (ThermoFisher Scientific)

1% BSA (Miltenyi Biotec)

0.2-1 U/uL Protector RNAse Inhibitor (Roche)

### Supplementary Figure 1: snRNAseq RNA sequencing quality control

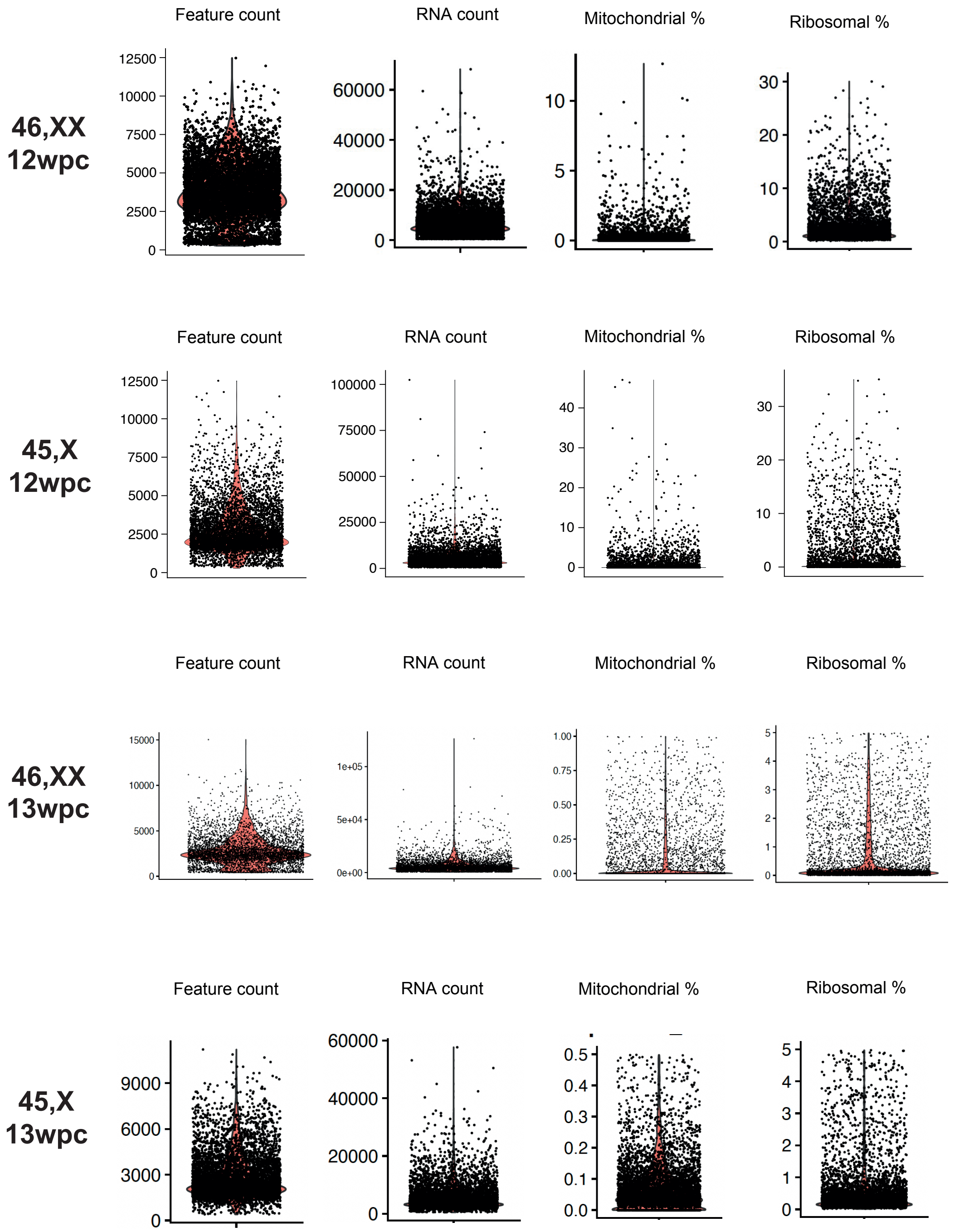


| | *13wpc 46,XX* | *13wpc 45,X* | *12wpc 46,XX* | *12wpc 46,X* |
|---|---|---|---|---|
| Raw cells | 5121 | 6164 | 7033 | 5792 |
| <1% mitochondrial counts | 4884 | 5849 | 6745 | 4972 |
| <5% ribosomal counts | 4732 | 5723 | 6180 | 4790 |
| Unique feature counts >400 | 4705 | 5723 | 6179 | 4790 |
| Doublets removed | 4489 | 5414 | 5802 | 4570 |
| **Final cell count** | **4489** | **5414** | **5802** | **4570** |

**Upper panel:** For each of the four samples included in the snRNAseq analysis, the feature count, RNA count, percent mitochondrial genes and percent ribosomal genes are shown.

**Lower panel (table):** The starting number of raw cells is shown for each ovary sample. Cells were removed from analysis if they had >1% mitochondrial counts, >5% ribosomal counts, unique feature counts <400, or were doublets. The final cell count for analysis is highlighted in bold.

#### Supplementary Figure 2: Integrated UMAPs for snRNAseq analysis

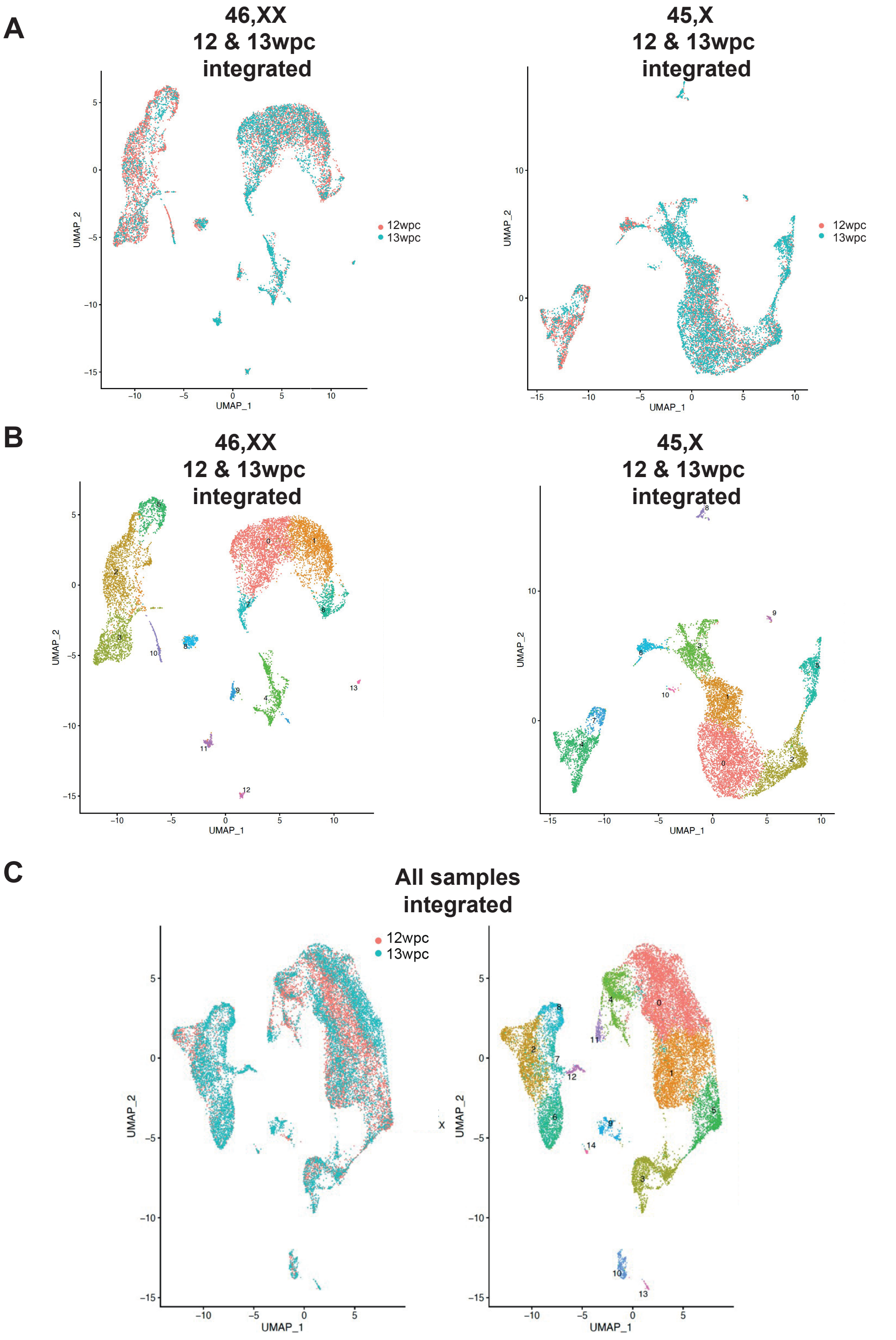


**Figure. UMAPs of integrated snRNAseq objects**

**A)** Integrated UMAPs of 46,XX ovaries (left) and 45,X ovaries (right). Red, 12wpc; Aquamarine, 13wpc; wpc, weeks post conception. **B)** Integrated UMAPs of 46,XX ovaries (left) and 45,X ovaries (right) after clustering (principal components 30, resolution 0.3). **C)** Integrated UMAP of all four samples split by age (left) and by clustering (right; principal components 30, resolution 0.3). Red, 12wpc; Aquamarine, 13wpc; wpc, weeks post conception.

**Supplementary Figure 3: Validation of key potentially relevant genes identified this study, using the Human Single-Cell Gonad Atlas (2022)**

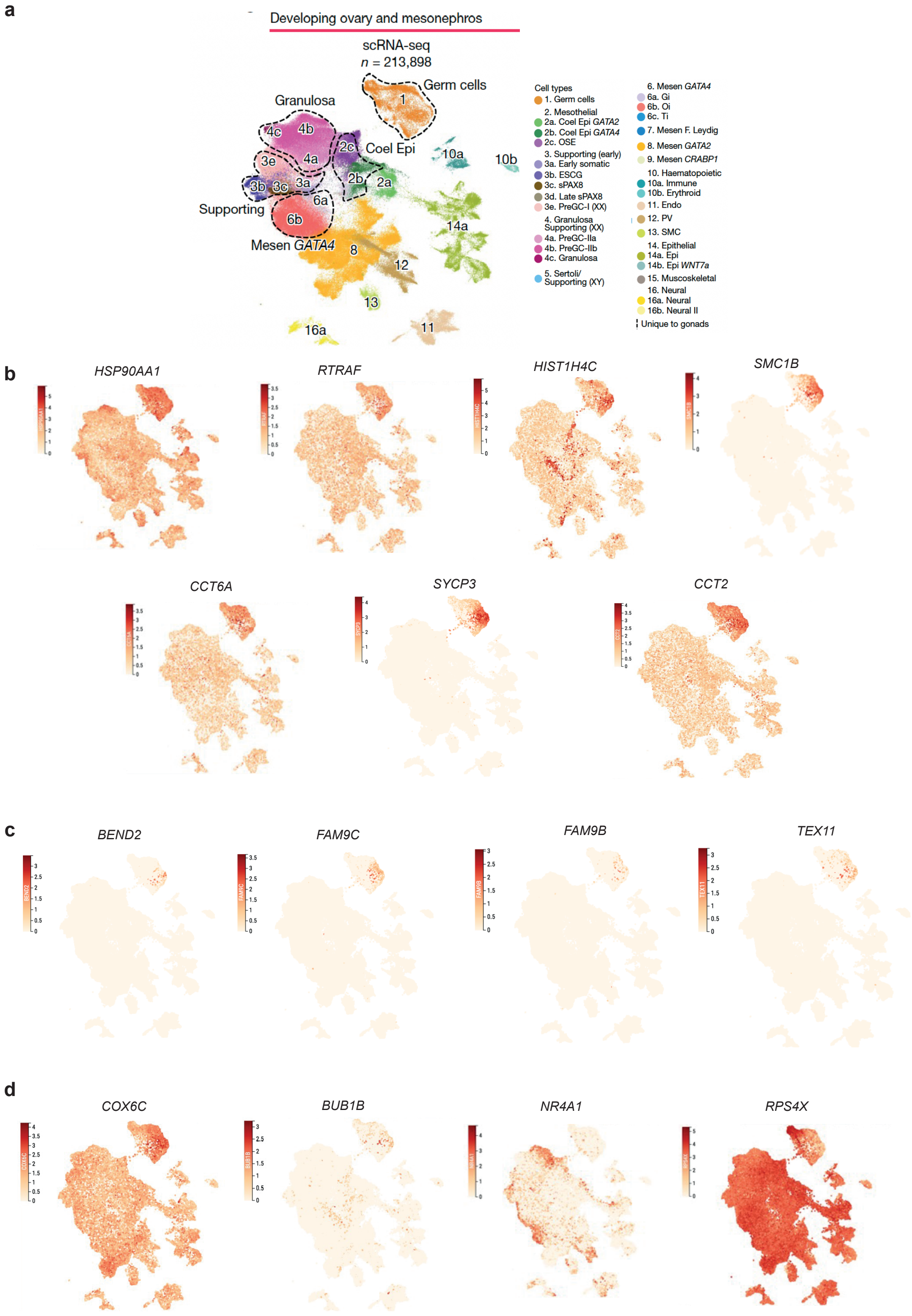


**A)** UMAP (uniform manifold approximation and projection) representation of previously published human fetal ovary single-cell data from Garcia-Alonso et al.

These data were generated by the Vento-Tormo group at the Wellcome Sanger Institute, Hinxton, UK and can be accessed using CZ CELLxGENE from https://www.reproductivecellatlas.org/gonads/human-main-male/ (Garcia-Alonso, L. et al. Single-cell roadmap of human gonadal development. Nature 607, 540–547, 2022; https://doi.org/10.1038/s41586-022-04918-4). This work is licensed under a Creative Commons Attribution-BY 4.0 International License (https://creativecommons.org/licenses/by/4.0/). **B)-D)** Key genes identified in this work are localised to the above Garcia-Alonso et al data for validation. B) Genes identified as highly expressed from the synaptic oogonia 46,XX-specific population; C) Key X chromosome/XCI escape genes discussed in the study; D) Genes identified as possibly involved in new mechanisms of ovarian insufficiency described in this study.

**Supplementary Figure 4. Bulk RNA sequencing experimental design**

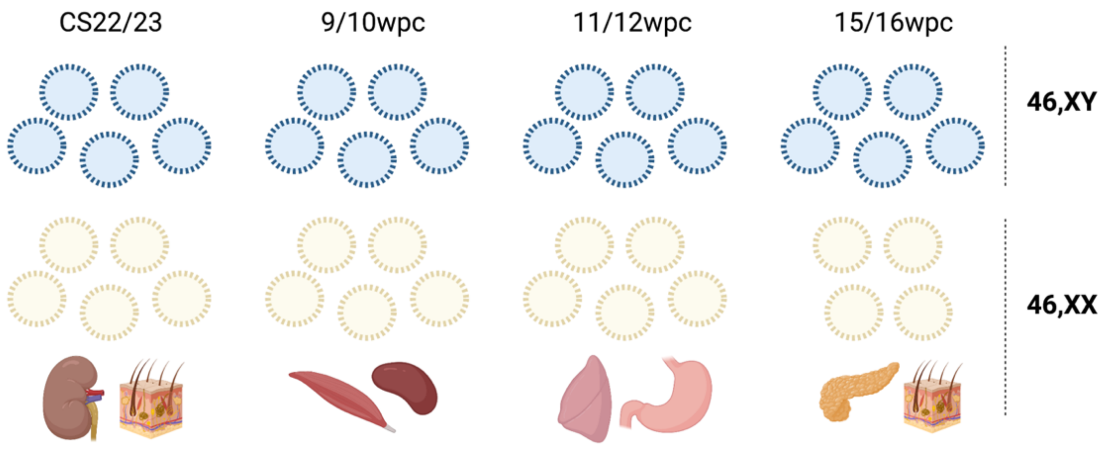


Testis (blue), ovary (yellow), and 46,XX control samples (kidney, skin, muscle, spleen, lung, stomach, pancreas) were collected for sequencing from each of four key developmental stages (CS22/23; 9/10wpc; 11/12wpc; 15/16wpc).

#### Supplementary Figure 5. Gene enrichment analysis of differentially expressed genes between 46,XX and 45,X clusters

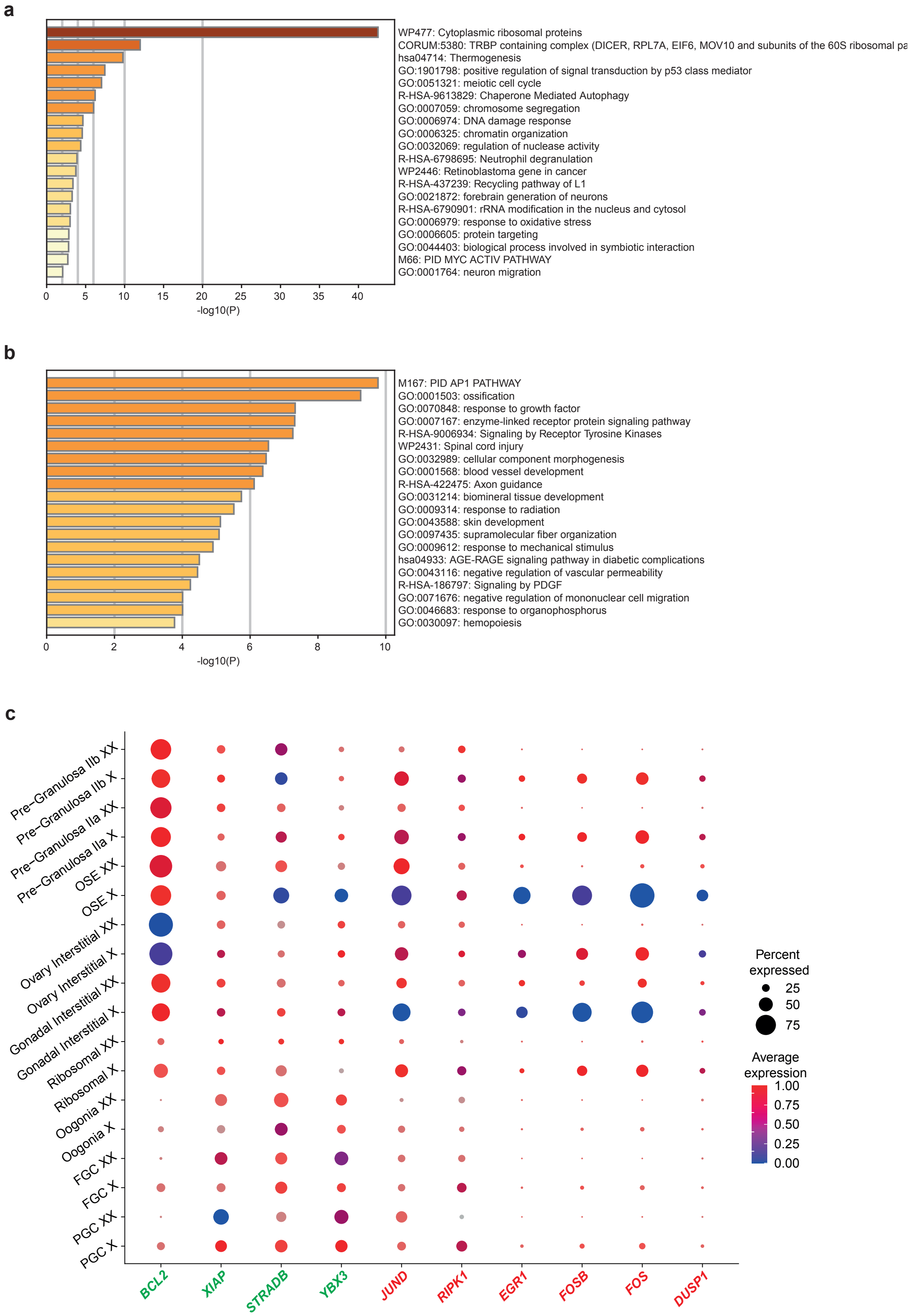

Gene enrichment analysis (Metascape) of **A)** genes positively differentially expressed in the 46,XX oogonia germ cell (PGC) cluster compared to the 45,X oogonia germ cell cluster; and **B)** genes positively differentially expressed in the somatic cell clusters of the 45,X ovary compared to 46,XX corresponding clusters (ovarian surface epithelium (OSE), interstitial cells, granulosa cells). **C)** Dot plot showing percentage and average expression of apoptosis-related genes across all individual cell clusters in 46,XX and 45,X ovaries. Genes in green have postulated anti-apoptotic roles; genes in red have proposed pro-apoptotic roles. FGC, fetal germ cell; PGC, primordial germ cell; OSE, ovarian surface epithelium.
